# Understanding the epidemiology and pathogenesis of *Mycobacterium tuberculosis* with non-redundant pangenome and population genetics

**DOI:** 10.1101/2024.10.25.620184

**Authors:** Yang Zhou, Richard Anthony, Shengfen Wang, Hui Xia, Xichao Ou, Bing Zhao, Yuanyuan Song, Yang Zheng, Ping He, Dongxin Liu, Yanlin Zhao, Dick van Soolingen

**Affiliations:** National Center for Tuberculosis Control and Prevention, Chinese Center for Disease Control and Prevention, No.155 Changbai road, Changping district, Beijing, China; Radboudumc Research Institute, Radboud University, Houtlaan 46525 XZ Nijmegen, The Netherlands; National Tuberculosis Reference Laboratory, Centre for Infectious Disease Control, National Institute for Public Health and the Environment (RIVM), Bilthoven, the Netherlands; National Institute for Public Health and the Environment, the Netherlands, 3720 BA Bilthoven, The Netherlands

**Keywords:** non-redundant pangenome, *Mycobacterium tuberculosis*, selective pressure, structural variation, evolution patterns

## Abstract

Tuberculosis is a major public health threat demanding more than one million lives every year. Many challenges exist to defeat this deadly infectious disease which address the importance of a thorough understanding of the biology of the causative agent *Mycobacterium tuberculosis* (MTB). We generated a non-redundant pangenome of 420 epidemic MTB strains from China. We estimate that MTB strains have a pangenome of 4,278 genes encoding 4,183 proteins, of which 3,438 of which are core genes. However, due to 99,694 interruptions in 2,447 coding genes, only 1,651 may be translated in all samples, which dramatically reduces the number of active core genes. Of these interruptions, 67,315 (67.52%) could be classified by various genetic variations detected by currently available tools, and more than half of them are due to structure variations, mostly small indels. We further describe differential evolutionary patterns of genes under the influences of selective pressure, population structure and background selection. While selective pressure is ubiquitous among these coding genes, evolutionary adaptations primarily occur in 1,313 genes. Genes located in the cell wall and membrane region are under the strongest selective pressure, while biological processes including regulation of transcription, translation and regulation of growth are under strongest background selection in MTB. The metabolism of fatty acids may be an outstanding example of evolutionary adaption for MTB under current selective pressure. This study provides a comprehensive view on the genetic diversity and evolution patterns of coding genes in MTB which may deepen our understanding of its epidemiology and pathogenicity.

## Introduction

Tuberculosis (TB) took 1.3 million lives in 2022 globally, making it the second deadliest infectious disease just after COVID-19.^1^ The World Health Organization (WHO) has set the goal to eliminate tuberculosis as a public health problem by 2035, which requires reducing TB incidence by 90% and mortality by 95% compared to 2015. New vaccines, better drugs and improved diagnostic tools are key advances needed to end the TB pandemic. However, only one ancient vaccine with limited efficacy, the so-called BCG developed from attenuated *M. bovis*, has been licensed for TB. Although two new drugs, bedaquiline and delamanid, have been added to the list of drugs available to treat TB in the past decades, strains extremely resistant even to these new drugs have already been observed. This stunted progress is partially due to the lack of incentives and underfunding, but also underlines our lack of understanding of the biological process and mechanisms relating to issues such as host-pathogen interaction and drug resistance development.

The first whole genome sequence of MTB strain H37Rv was published in 1998 and has been used as the backbone for mapping short reads from high-throughput sequencing platforms to identify genetic variations, primarily single nucleotide polymorphisms (SNPs) and small indels described in most studies.^2^ This information can be used to infer the phylogeny and evolution of epidemic strains, to monitor recent transmission and outbreaks, and to predict drug resistance phenotypes.^3–6^ These milestone findings suggest that SNPs and small indels indeed correspond to a large proportion of the observed diversities in clinical MTB isolates. However, the limitation of this pipeline is also obvious since genetic variations in the regions not presented in the reference genome are not included. Comparative genome analysis has already shown that large sequence polymorphisms (LSPs) between strains are not only related to the genealogical delineation, such as Lineage 2 of MTB defined by RD105, but are also associated with important biological processes.^7,8^ For example, two genes *mmpS6* and *mmpL6* are both deleted or truncated in “modern” MTB strains due to the deletion of TbD1 region, which may facilitate the successful expansion of modern strains globally through a hypoxia-induced copper response.^9^ Another region of difference, RD1, harbors several open reading frames (ORFs) encoding ESX-1 secretion system proteins including CFP-10 and ESAT-6 which are crucial to virulence and phagosomal escape.^10,11^ The RD1 deletion in *M. bovis* is related to the attenuation of the BCG vaccine.^12^ Besides these LSPs which result in complete deletions of genes, other types of structural variations (SVs), such as IS*6110* insertions in coding sequences, partial deletions of genes by large deletions (>100 bp) or interruptions by large insertions, are also missed by the standard pipeline and may have important consequences related to important biological processes.^11,13,14^ By focusing only on SNPs and small indels, it’s assumed that at least a certain proportion of genetic diversity information in clinical strains of MTB is missed, which leads to the development of a more comprehensive approach to study genomic data.

Accumulation of high through-put sequencing data has allowed the development of a new study field: pangenome analysis.^15,16^ The term pangenome refers to all the genetic content of a given dataset, usually a species. The shared genomic content or genes in all genomes are referred to as core genes; other genes presented in several but not all genomes are called dispensable or accessory genes which may be introduced by horizontal gene transfer, gene duplication/deletion or other SVs. Accessory genes are supposed to account for much of the diversity among clinical strains of microorganisms.^17^ This is especially true for species with a large number of accessory genes, such as *E. coli*, which has high genome plasticity and a very large gene pool.^18^ MTB is known to have a conserved genome with no plasmids detected and horizontal gene transfer is very rare in modern strains if it exists at all. Thus, it’s questionable how much of the diversity observed in clinical practice could be explained by the presence/absence of accessory genes. The drawback of the presence/absence approach is that the variations within genes, such as nonsense mutations and frame shift mutations, are ignored, which might contribute to the presence/absence at protein level.^19^ Thus, for a microorganism with a conserved genome such as MTB, a combined approach by integrating the pangenome and within genes variation information is possible.

In this study, we developed a pipeline to generate a non-redundant pangenome of MTB addressing the problems outlined above from the available short read sequencing data.^20^ Based on the pangenome matrix and variations detected with dedicated tools, we described the genetic variation at nucleotide, gene and protein levels. Based on this comprehensive profile of genetic diversity in the pangenome, we characterized the evolutionary pattern of all coding genes under the influence of selective pressure, population structure and background selection, which reflects the complex interactions between these factors. Gene ontology analysis of genes showing different evolutionary patterns has revealed some primary characteristics of the MTB biology and the interactions between MTB and the environment.

## Method

### Sampling and WGS

The 420 clinical MTB isolates used in this study were selected from the first national drug resistance baseline survey in China in 2008 (available in **PRJNA573798**, at https://www.ncbi.nlm.nih.gov/bioproject/PRJNA573798/). Selection criteria were based on drug resistance patterns, geographic location of isolation, and genotypes such as spoligotype and MIRU-VNTR profile. The phylogenetic and evolutionary analysis based on whole genome sequencing were published previously.^20^ Briefly, all samples were sequenced on an Illumina platform using single end or paired end library. After filtering out repeat regions, PE/PPE_PGRS genes and insertion sequences, phylogenetic trees were constructed with MEGA using a maximum likelihood algorithm.^21^

### Assembly and annotation

The same sequencing data was used to produce the assembly of each strain using the program Spades with k-mer size auto-detected and coverage cut-off set to off and trying to reduce the number of mismatches and short indels.^22^ The resulting contigs were first blasted against the NCBI nucleotide database to remove possible contamination from the previous steps. All blast hits of e- value >1e-5 were excluded. Preserved contigs were excluded if aligned to species other than MTBC. The quality of assemblies was evaluated with QUAST.^23^ All contigs longer than 200 bp that passed the contamination filtering were annotated by Prokka ^24^ to predict coding sequences (CDSs) with default parameters. The produced gbk file was used to search for homologues genes.

### Constructing non-redundant pangenome

Two approaches are combined to get the full-set of non-redundant pangenome: CDS prediction and blastn. CDSs predicted by Prokka were clustered by GET_HOMOLOGUES using COG algorithm. ^25^ When searching for clusters, the max e-value for blast was set to 0.01, the max intergenic size was set to 1,000 bp, minimum coverage was using the default 75%. At least one sequence was required for a cluster to be called. Only CDSs were considered for clustering. Clusters which were clustered or overlapped with Rv by >=60 bp were considered as redundant to Rv and were removed from candidate new genes list.^26^ The left candidate new-/non-Rv genes went through self-overlapping filter by length to keep the longer CDSs if the overlap exceeds 60 bp.

Because Prokka can only predict valid ORFs, for those with interruptions such as frameshift mutations or other structure mutations, these genes are not predicted in the output or predicted with alternative ORFs, but could provide information on phylogeny and evolution. To further detect this part of information missed by prediction software, we applied blastn using consensus sequences of each cluster as the query sequence. Consensus sequences of the new candidate genes and sequences of all 3,906 CDSs and 112 other genetic features from H37Rv genome, including pseudogenes (30), tRNAs (45), ncRNA (20), rRNA (3), misc RNA (2), fragment of putative small regulatory RNA (10), and two other feature genes, were blasted to all contigs of 420 samples and the H37Rv genome. Blast hits with e-value < 1e-5, identity > 90% and query coverage > 30% (for Rv genes) / > 90% (new candidate genes) were kept. Blastn results went through redundant checking using synteny and overlapping information. When two blast hits for the same gene in one sample have no overlap of their locations, they would be considered as gene duplications and only represented one homologues gene.

Finally, candidate new genes in complementary sequences to H37Rv genome were manually checked and filtered according to expert opinion by manual checking the similarity of complementary sequences and locations of the candidate new genes in the complementary sequences. All contigs were blasted to the H37Rv genome sequence and blast hits with e-value < 1e-5 and identity > 90% with the highest score were kept. Complementary sequences, absent in the H37Rv genome, were extracted based on blast results. All complementary sequences longer than 100 bp were compared to each other and those sharing more than 90% identity and more than 95% query coverage with e-value < 1e-5 were considered as homologous sequences. This step could filter alternative ORFs predicted in different samples that were not filtered out in previous steps.

### Pangenome composition

The presence or absence of an individual gene was analyzed at different levels separately. At DNA level, if blastn identified a non-redundant best hit for a gene, it was considered as the presence of this gene. At protein level, only when this gene sequence had a valid ORF it was considered as present. Conversion of gene sequence to protein sequence was done with Biopython package.^27^ Non-synonymous mutations in start/stop codon resulting in loss of start/stop codon, gain of extra stop codon, frameshift mutations, and IS*6110* insertions in CDSs were considered as interruptions of translation. In addition, if > 90% of the gene sequence was deleted or not detected by blastn, it was considered as complete deletion.

### Detection of structural variations

Eleven open source software packages were used to call insertions and deletions, including assemblytics^28^, bcftools^29^, breakdancer^30^, delly^31^, lumpy-smoove^32^, minimap2^33^, svaba^34^, softsv^35^, tiddit^36^, unimap^37^, and wham^38^. Called variants were first filtered based on the parameters with each method such as number of supporting reads. Inversions were not analyzed in this study. Deletions and insertions > 100 kb were also filtered without further check. Variants detected for each sample were merged by SURVIVOR only if supported by more than one method (for deletions > 1,000 bp, more than four methods were required) with restrain to the same variation type and breakpoints within 100 bp.^39^

The pangenome matrix at gene level from composition analysis was compared with the results from SURVIVOR. Deletions >1,000 bp without corresponding gene deletion in the pangenome matrix were checked manually with IgV.^40^ The gene deletions in the pangenome matrix that were not detected by the 11 methods were checked by viewing in IgV and added to the dataset if confirmed. IS*6110* insertions were detected in paired end samples with ISMapper.^41^

### Genetic diversity, selective pressures, positive selection and population stratification

Tajima’s D ^42^ and nucleotide diversity π ^43^ for each coding gene were calculated using Dnasp.^44^ Interrupted gene copies were excluded before calculating the statistics under the assumption that these sequences evolve faster than valid ORFs. Significance for Tajima’s D was given by the table in the original paper by Tajima.^42^ Homoplasy was detected by homoplasyFinder using the default parameters.^45^ Homoplasy in SNPs were only detected in valid CDSs. For each gene tree used to find homoplasy SNPs in the specific gene, the phylogeny tree produced previously was re-used and genomes with interrupted CDSs were removed using python package ete.^20,46^ SVs resulting in interruptions of the same CDS were considered as homoplasy ignoring the type and the position information. Interquartile range (IQR) for homoplasy events was calculated by substracting the 25^th^ percentile from the 75^th^ percentile. High homoplasy level was defined as more than the sum of IQR and the 75^th^ percentile. Fixation index Fst was calculated using command basic.stats from R package Hierfstat according to the equation 7.38-7.43 in page 164-5 of Nei (1987).^47,48^ All negative Fsts were considered as zero. Significance for positive Fsts were calculated by permutating all samples among all groups for 1,000 times and counting the number of Fsts greater than or equal to the observed Fst using the same R package Hierfstat.

### Gene ontology and overrepresentation analysis

Coding genes under different evolutionary patterns were subjected to gene set over-presentation analysis using the online tools provided by PANTHER (version 19.0 based on GO database published in Jan 17, 2024).^49^ Bonferroni correction and Fisher exact test were implemented to find over-presented GO items as compared to the standard GO annotation database. Adjusted P-value < 0.05 was considered as significant. Only the most specific GO items were retrieved.

## Results

### *M. tuberculosis* has a conserved and closed pangenome

The data sample used in this study has been analyzed in a previous study which described the phylogeny and evolution of the epidemic MTB strains in China.^20^ This sample set contained 344 Beijing family strains, 14 Lineage 4.2 strains, 15 Lineage 4.4 strains, 40 Lineage 4.5 strains, 6 Lineage 3 strains and one Lineage 1 strain, which were representative of the epidemic MTB strains in China and these strains were also prevalent in other regions globally.

The number of coding genes annotated for each genome varied from 4,167 to 4,337 (median = 4194). The initial pangenome set automatically predicted by software consisted of 10,097 coding gene clusters, which included 3,754 singleton clusters. More than two third (1,167,634; 68.38%) of the predicted CDSs were identical to the blast results. Besides Rv gene sequences, 66,412 sequences in 436 clusters remained after removing redundant sequences from CDSs predicted by prokka. After blasting consensus sequences of these 436 clusters to all contigs and repeating the process of removing redundance, 288 clusters were left. At this step, one cluster was removed because there were only two sequences of very different lengths in this cluster and no consensus sequence could be produced; the other 147 clusters were removed because some CDSs in these clusters were missed by prokka due to interruptions in the ORF which were detected by blastn and overlapped with other CDSs for >= 60 bp.

Among the remaining 288 clusters, 20 clusters harbored repeat sequences or were in insertion sequences and were removed from the final pangenome. There were 175 clusters which had homologues sequences in H37Rv genome and were considered as new Rv genes. Among the other 93 clusters, 22 clusters were in previously reported LSPs, such as RvD1 (four clusters), RvD2 (4), RvD5(4), and TbD1 (1), and some less known LSPs, such as the RvD4494 (inserted at 2,219,418, six clusters). Another 40 clusters were in complementary sequences adjacent to PPE/PE_PGRS genes or IS*6110*/IS*1532* insertion sequences or both. Seven clusters were in complementary sequences adjacent to other highly polymorphic regions, such as *Rv2082*. These 69 clusters were not detected in H37Rv genomes and were considered as non-Rv genes. Another 16 clusters were not in complementary sequences to H37Rv, but were interrupted by various kinds of SVs, such as *15427_dxs* interrupted by IS*6110*-15 insertion. These 16 clusters were also considered as non-Rv genes, which resulted in a total of 85 non-Rv genes. The eight clusters left were excluded from the pangenome because of overlapping >= 60 bp in H37Rv (though in some genomes < 60 bp) or being in low quality short contigs (about 300 bp). Finally, 260 new genes were added to the final pangenome in addition to the 4,018 genes/features in H37Rv genome, including 175 new Rv genes and 85 non-Rv genes.

The final non-redundant pangenome constituted 4,278 genes, 4,183 of which were protein coding genes. Besides the 3,906 CDSs and 175 new Rv genes in the H37Rv genome, another 17 pseudo Rv genes interrupted in H37Rv might produce translation products in other genomes, so the total coding genes in H37Rv were 4,098. (**Fig. 1**)

**Figure 1.**
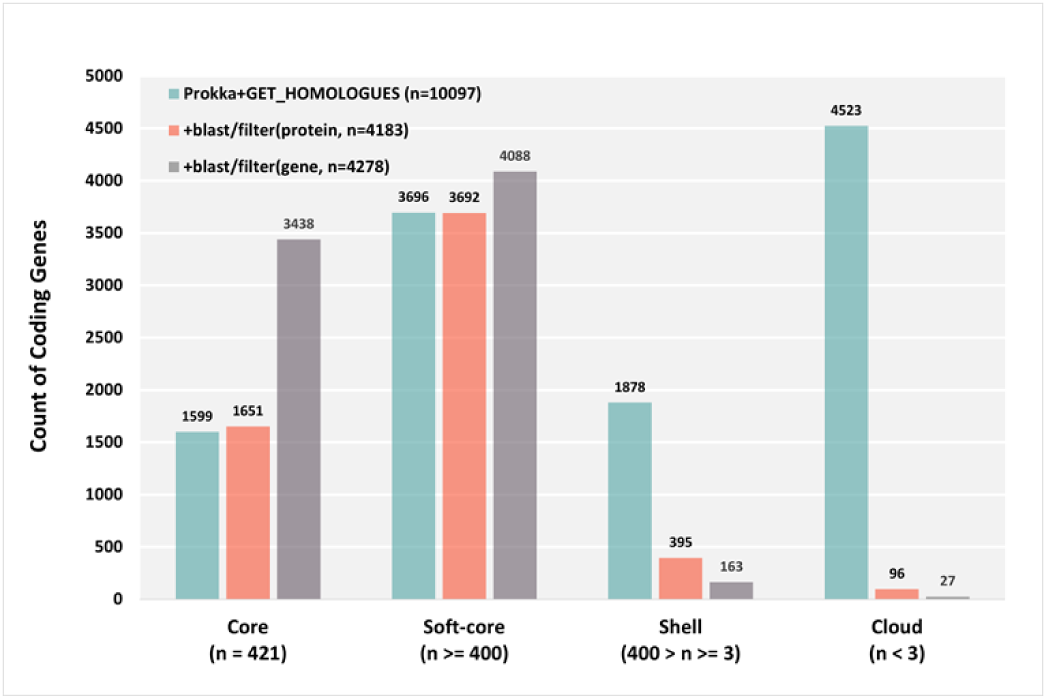
Pangenome composition. Pangenome composition at different levels and with different methods.

### Genetic diversity in the pangenome

Among the 4,183 protein coding genes in this pangenome, 3,438 genes were presented in all genomes, but only 1,651 genes had valid ORFs in all 421 genomes. There are 99,694 interruptions, defined as no protein sequence produced by *in silico* translation, in 2,447 coding genes. We combined pangenome matrix and genetic variations detected by dedicated tools to see how genetic variations affected the presence or absence at different levels in the pangenome. Because most of the available tools to detect SVs and SNPs with short reads sequencing data require a linear reference genome, genetic variations such as SNPs and SVs were only detected for the 4,098 coding genes in H37Rv genome.

Software for SVs detected 76,671 variants in the 4,098 coding genes in total, including 49,655 deletions, 26,700 insertions and 316 inversions. Inversions were not included in the following analysis. For the 325 paired-end samples, 1,860 IS*6110* insertions at 411 positions in 311 samples were detected. There were 16,151 high impact point mutations in start/stop codons, such as those causing loss of a start/stop codon or gain of an extra stop codon, detected in 1,007 genes for 421 samples using the standard pipeline.

There were 23,221 gene copies which might be completely deleted and could not be detected by blastn, of which 50.40% (11,704) could be attributed to large deletions. High impact point mutations in start/stop codons alone caused 12.65% (12,615) interruptions in CDSs, while other SVs corresponded to another 46.26% (42,132) interruptions in CDSs besides complete deletions. Small indels (<100 bp) contributed the largest part of interruptions (31.22%). Together, SVs and high impact point mutation in start/stop codons explained 67,315 (67.52%) interruptions in the pangenome in total, which left 32.48% of the interruptions unclassified. (**Table 1**)

**Table 1.** Genetic variations with interruptions in coding genes.

| Category | Genetic Variation | Interruptions | % |
| --- | --- | --- | --- |
| **SV** | Indel | 31,123 | 31.22% |
|  | Complete deletion | 11,682 | 11.72% |
|  | Large deletion | 5,152 | 5.17% |
|  | Large insertion | 1,463 | 1.47% |
|  | IS6110 | 1,735 | 1.74% |
|  | complex SV | 2,659 | 2.67% |
|  | **sub total** | **53,814** | **53.98%** |
| **SNP** | stop gained | 6,505 | 6.52% |
|  | stop lost | 1,961 | 1.97% |
|  | start lost | 4,148 | 4.16% |
|  | complex SNP | 1 | 0.00% |
|  | **sub total** | **12,615** | **12.65%** |
| **Complex** |  | **886** | **0.89%** |
| **Interruption** | **classified** | **67,315** | **67.52%** |
|  | **un-classified** | **32,379** | **32.48%** |
|  | **total** | **99,694** |  |

A total of 32,379 un-classified interruptions were observed in 1,328 genes. There were 253 genes having more than 21 (5% of total samples) un-classified interruptions. The un-classified interruptions in these 253 genes corresponded to 88.29% (28,587) of all un-classified interruptions detected in the 2,447 coding genes. A previous study identified 516 genes in blind spots in the H37Rv genome in Illumina sequencing, 191 of which intersected with these 253 genes.^50^ There were 23,940 (71.44%) un-classified interruptions in the 516 genes in the blind spots, 23,097 of which were in the 191 genes in the intersection, leaving the other 325 genes in blind spots with only 843 un-classified interruptions, indicating some previously identified blind spots might have produced high quality results in our dataset. Beyond the 191 genes, there were 9,282 interruptions un-classified in 1,137 genes. There were 33 samples each with > 50 un-classified interruptions of in total 3,635 (39.16%) un-classified interruptions, indicating these samples might have yielded lower sequencing quality and more errors in assembly, but this bias was less obvious as compared to genes in blind spots. After excluding the 253 low quality genes, there were 46,024 interruptions in 2,194 genes and only 8.24% (3,792) of the interruptions were not classified by genetic variations.

Some of the genetic variations were phylogenetically specific, such as TbD1 present only in Lineage 1, which harbored two ORFs *mmpS6* and *mmpL6*, and RvD2 absent in Lineage 2 and Lineage 3 which harbored four ORFs including *pimC*, *suoX*, *15952_hypothetical_protein* and *15953_transmembrane_transp*. We have identified several other LSPs which were lineage specific. A fragment of 3,138 bp (inserted between 1,414,558-1,415,891 in H37Rv) was deleted in Lineage 2 and Lineage 4 strains harbored two ORFs. This fragment of complementary sequence could be fully matched to other MTBC genomes in NCBI database but not to the H37Rv genome, possibly another LSP yet to be reported. Another LSP of 4,494 bp (inserted to H37Rv at 2,219,418) was (partially) deleted in all Lineage 4 strains but present in all other genotypes, which contained six ORFs and has been reported in a previous study.^7^ In the other hand, some LSPs were found in highly polymorphic regions where multiple overlapping SVs were identified in different genomes. For example, RD3, which harbored sixteen genes (including one gene of low quality) were deleted in Lineage 2 strains; however, RD3 was also deleted in 21 Lineage 4 strains. In the region where RD152 was located, multiple deletions were observed in Lineage 2 and Lineage 4 strains but with different borders: eight genes (including two of low quality) were absent in Lineage 2 strains due to the deletion from 1,986,638 to 1,998,622; six genes were deleted in these 21 Lineage 4 strains due to the deletion from 1,987,702 to 1,998,657. In Lineage 2 strains, 40 genes were absent due to LSPs, including 14 phage related genes and five CRISPR associated genes.

High impact SNPs or other types of SVs could also result in interruptions of genes in specific lineages. For example, *Rv0197* was interrupted by an insertion of 2 bp at position 234,496 in Lineage 2 strains but also in all other strains except H37Rv; *ephF* was interrupted by the deletion of one nucleotide at site 162,152 in Lineage 2 strains; *Rv0061c* was interrupted by a point mutation from G to T at position 65,150 which resulted in the gain of an extra stop codon in Lineage 2 strains. *15427_dxs* was interrupted by IS*6110*-15 insertion in H37Rv; in our dataset, no other genomes had this IS*6110* insertion but at least 351 genomes had an insertion of 5 bp in this gene, so only 66 genomes (all Lineage 4) had valid ORFs of this gene in our dataset. In total, interruptions caused by high impact SNPs and SVs other than LSPs resulted in the absence of 40 genes in Lineage 2 strains in our dataset.

### Landscape of genetic diversity shaped by selective pressures, population structure and background selection

To investigate how genes evolve under selective pressures, we calculated Tajima’s D and nucleotide diversity π and detected homoplasy events in coding genes. The 253 coding genes that might have inferior sequencing quality and the 85 non-Rv genes that were prone to errors with short reads sequencing were excluded for this analysis, thus 3,845 genes were analyzed.

Nucleotide diversity was calculated for 3,818 coding genes with >1 valid CDS which varied between 0 to 7.09e-3. The average nucleotide diversity was 1.51e-4 in this pangenome. There were 957 genes of higher nucleotide diversity than the average and 2,861 genes of lower nucleotide diversity. (**Fig. 2**)

**Figure 2.**
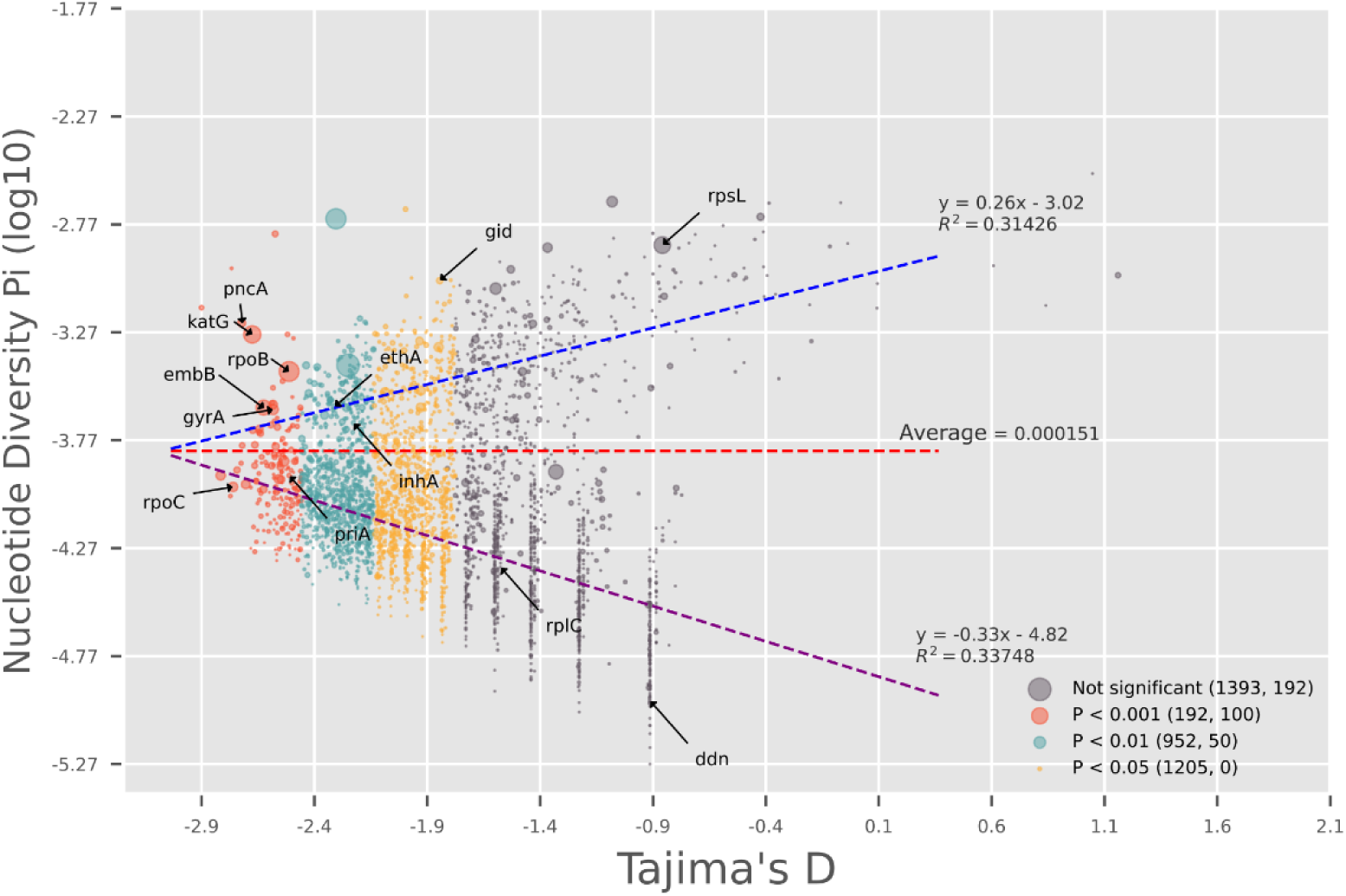
S**e**lective **pressure, nucleotide diversity, and homoplasy in the pangenome (n=3,742).** Only high quality Rv genes with > 3 valid CDSs and at least one segregating nucleotide site are shown. Each circle represents a gene. x axis is Tajima’s D, and y axis is nucleotide diversity π. Count of homoplasy events are shown by the size of the circles. The colors show the different levels of selective signals. The first number in the parenthesis is the number of samples at that level of selective pressure, and the second number shows the corresponding number of counts of homoplasy events as shown by corresponding circle size. The red line shows the average nucleotide diversity across the pangenome. The blue line shows the predicted π by linear regression model for genes above the average genetic diversity, and the purple line shows the predicted π below the average genetic diversity.

Tajima’s D was obtained for 3,742 coding genes with > 3 valid CDSs and at least one segregating nucleotide site, 2,349 of which have shown significant signal of selective pressure (adjusted P-value < 0.05) and 192 showed strong signal of selective pressure (adjusted P-value < 0.001). (**Fig. 2**) Among the 2,353 genes with information of cellular component, 141 (7.54%) of the 1,870 genes in cell wall/membrane regions (GO:0009274, GO:0016020, GO:0005886, and GO:0042597) were under strong selective pressure, while only 18 (3.73%) of 483 genes not in cell wall/membrane were under strong selective pressure (P-value = 0.004). Most of the drug resistance related genes including *gyrA*, *priA*, *rpoB*, *rpoC*, *katG*, *pncA*, and *embB*, showed signals of selective pressure, except *rpsL, rplC* and *ddn*. While *rplC* and *ddn* were conserved in sequences (π < 1e-5), *rpsL* showed high nucleotide diversity (π = 0.001356). The genetic variation in *rpsL* in our dataset was constrained to four nucleotide loci and mostly in codon 43 (72.92% of K, 0.24% of M, and 26.37% of R). On the contrary, in other drug resistance related genes under selective pressures, such as in *katG*, many low frequency alleles were observed. (**Fig.3**) There were 71 genes with > 3 valid CDSs having no segregating sites thus no genetic variation detected, 40 of which were core genes.

**Figure 3.**
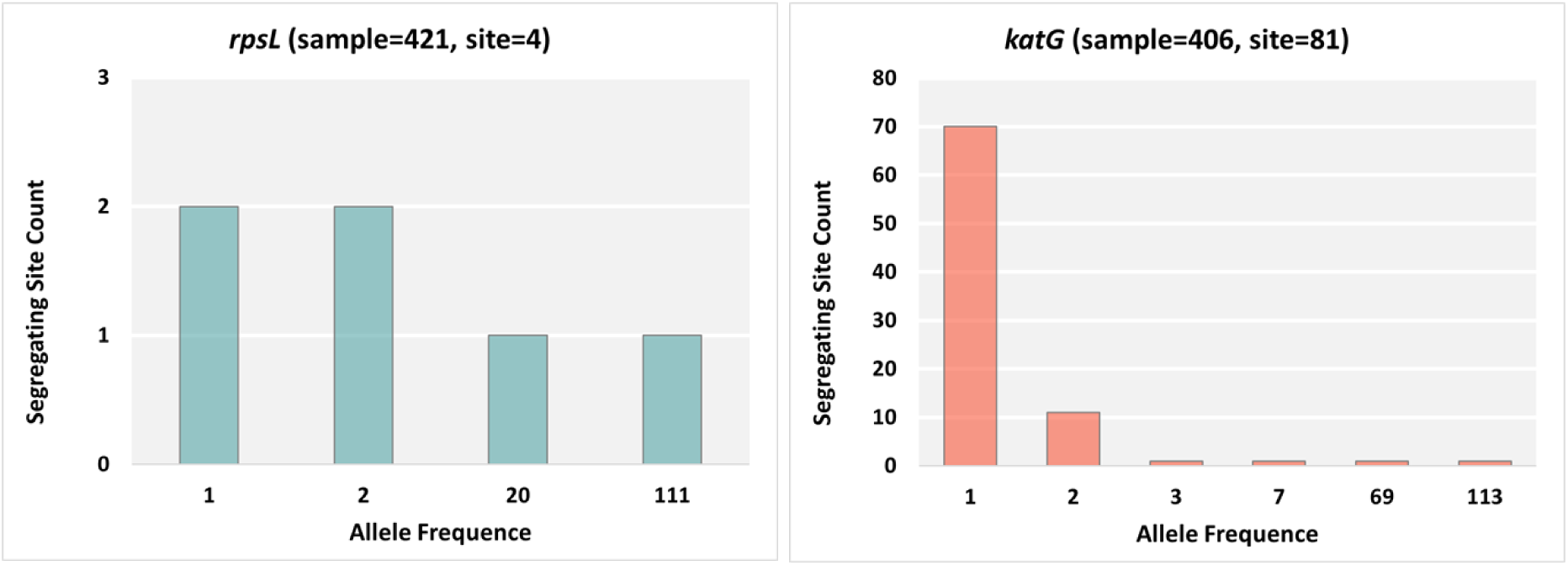
Folded (ranging from 0 to 0.5) minor allele frequency spectrum for *rpsL* and *katG*.

Selective pressure had a complicated impact on nucleotide diversity. For genes of high nucleotide diversity, this phenomenon was negatively correlated with the strength of selective pressures (or positively correlated with the value of Tajima’s D); for genes with low nucleotide diversity, this phenomenon was positively correlated with the strength of selective pressure. (**Fig. 2**)

Frequent deletions or high impact point mutations were observed in some regions in the pangenome, indicating homoplasy in the evolutionary development in these regions and genes. (**Fig. 4**) We have identified 6,695 homoplasy events in 1,745 coding genes including 3,159 SVs/high impact point mutation homoplasy events and 3,536 homoplasy events with SNPs. There were 129 genes showing higher level of homoplasy (>= 9 homoplasy events). The most significant homoplasy SNPs in our dataset was in codon 315 in *katG*, which was estimated to have mutated 92 times. (**Fig.5**). For homoplasy caused by SVs, interruptions of *esxR* were observed 82 times due to different deletions and insertions, which was the most frequently observed interruption by SVs. (**Fig. 4**) Disruption of host anatomical structure (GO:0141060) was over-presented by five genes, four of which had phospholipase activity (GO:0004620) including *plcA*, *plcB*, *plcC* and *Rv08 88*. In addition, all three proteins of sphingomyelin phosphodiesterase activity (GO:0004767) and six proteins of phosphopantetheine binding (GO:0031177) activity were also over-presented in this group. (**Table 2**)

**Figure 4.**
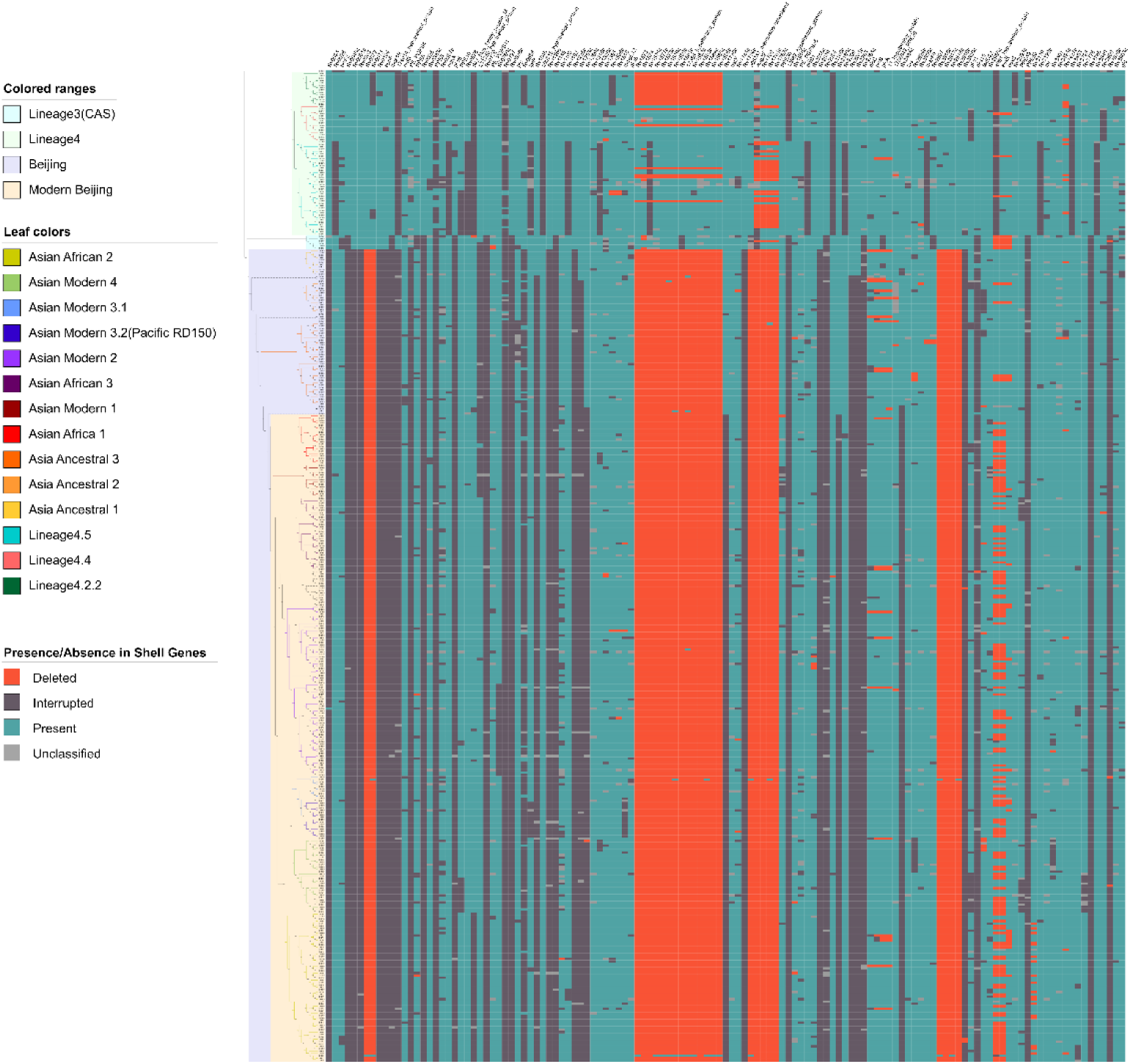
Genetic variations in shell genes. 127 shell genes in the pangenome with information of presence and absence at protein level in each sample. Low quality genes and non-Rv genes were not presented. The phylogeny tree on the left side was constructed in our previous report using 33,220 SNPs.^20^ Bright red boxes represent genes completely (>90% of length) deleted in that genome; dark grey boxes represent genes presenting in the genome but were interrupted by SVs or high impact SNPs; blue boxes represent genes with valid ORFs; light grey represent genes with interruptions due to unclassified factors.

**Figure 5.**
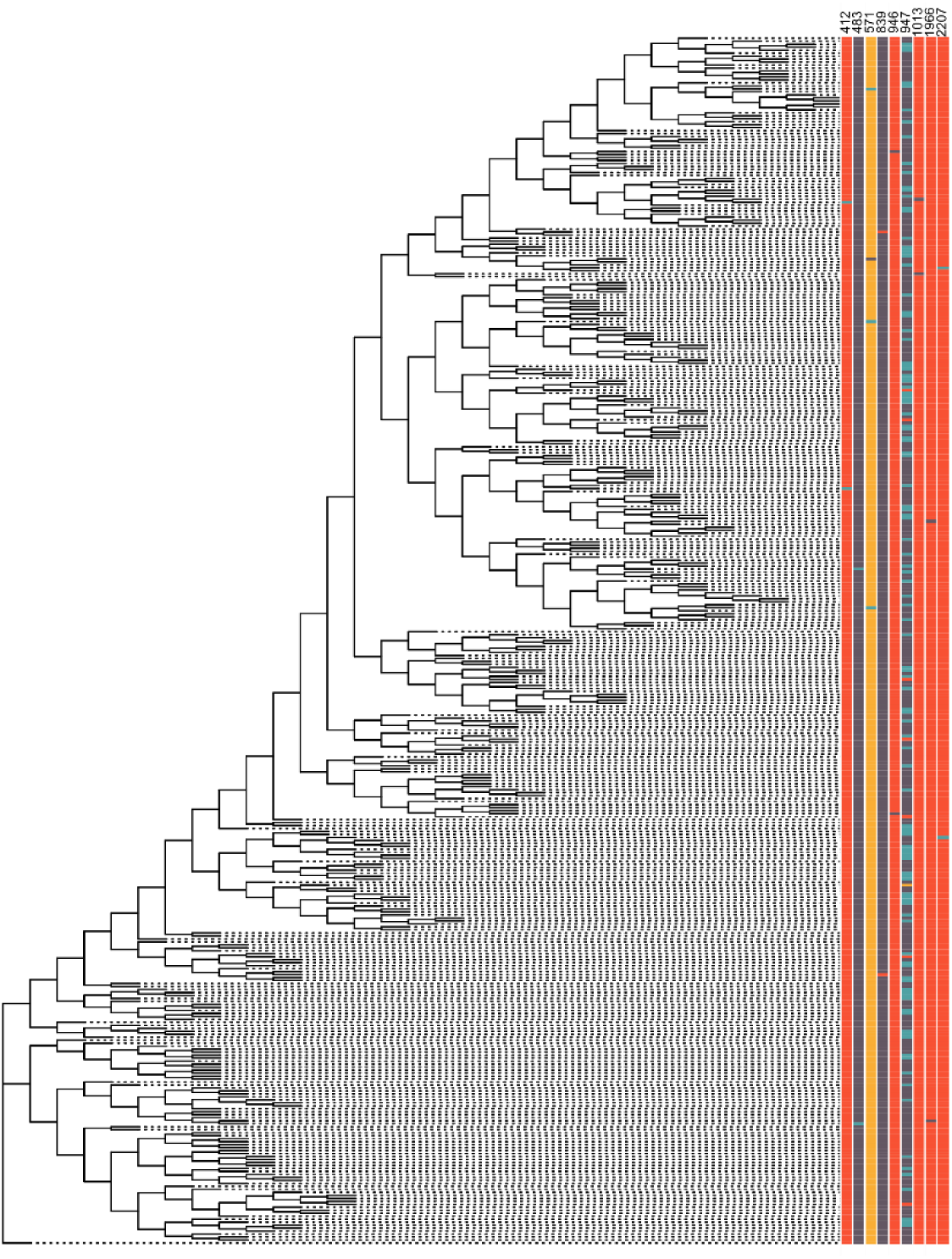
H**o**moplastic **SNPs in *katG*.** Phylogeny tree shows 404 genomes with valid *katG* ORFs. Nucleotide site 947 corresponds to 944 in H37Rv due to a 3 bp insertion in some genomes upstream of this position. Strips show the sites with homoplasy with nucleotide in different colors: A=red, C=blue, G=grey, T=orange.

**Table 2.**
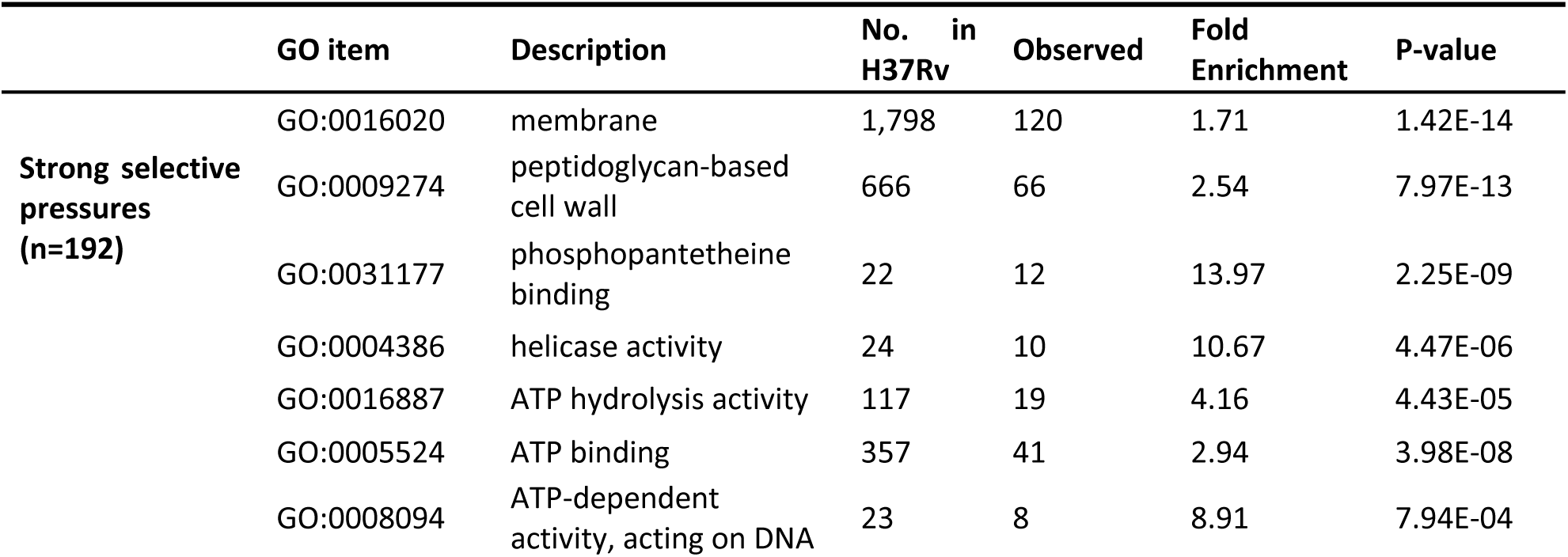
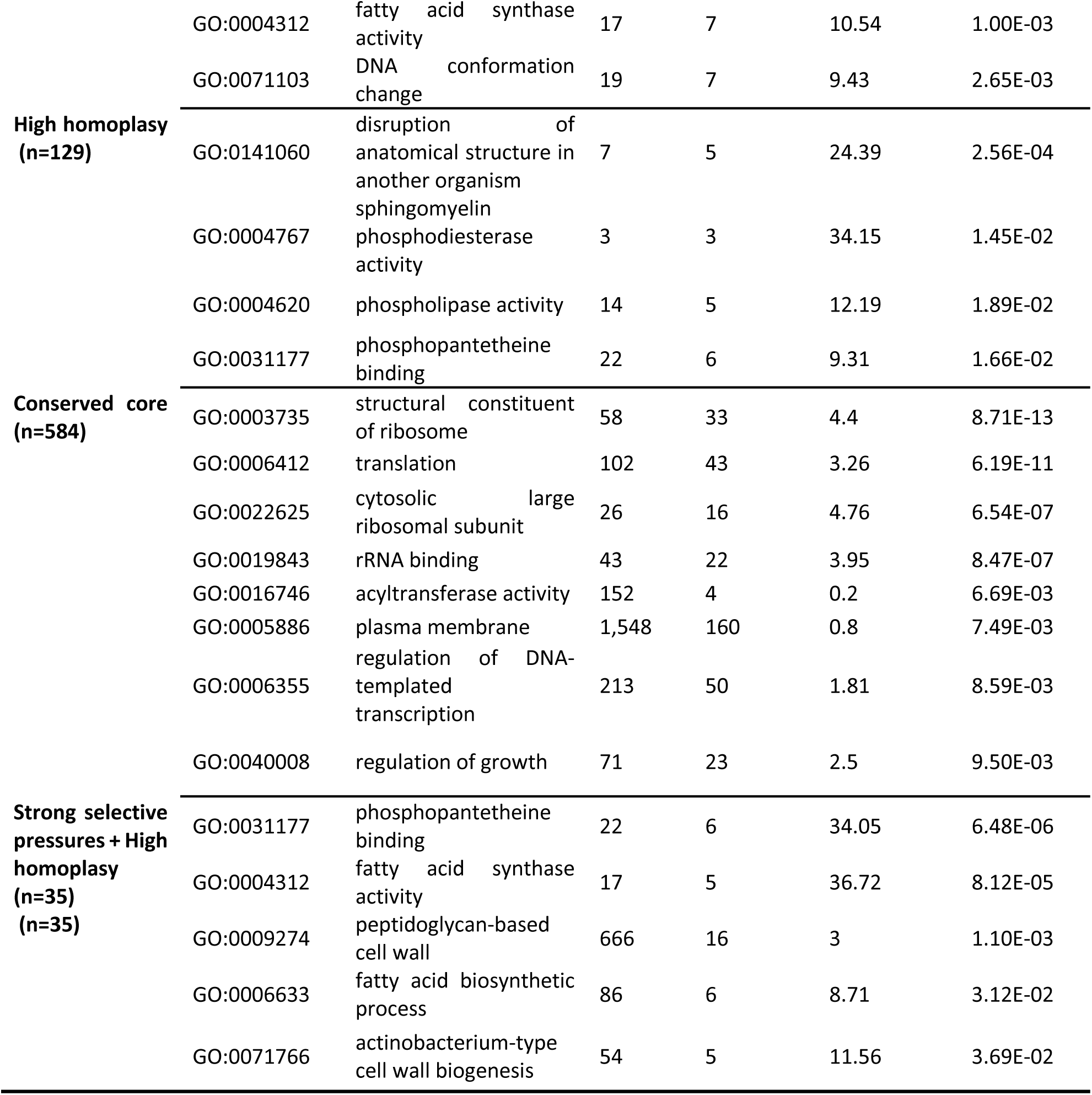
Over-presented gene categories.

Under selective pressure, 1,310 genes have shown signals of positive selection as indicated by homoplasy. The level of homoplasy was positively correlated to the strength of selective pressures in these genes. (**Table 3**) For the 192 genes under strong selective pressure, 155 (80.73%) genes showed homoplasy, and the average number of homoplasy events was 8.63; for the 1,393 gene showing no signal of selective pressures, only 406 (29.15%) genes showed homoplasy and the average number of homoplasy events was 1.04. There were 35 genes showing high levels of homoplasy under strong selective pressure. Besides six genes related to drug resistance, fatty acid biosynthetic process (GO:0006633) was over-presented by six genes in this group including five genes with fatty acid synthase activity (GO:0004312). (**Table 2**)

**Table 3.**
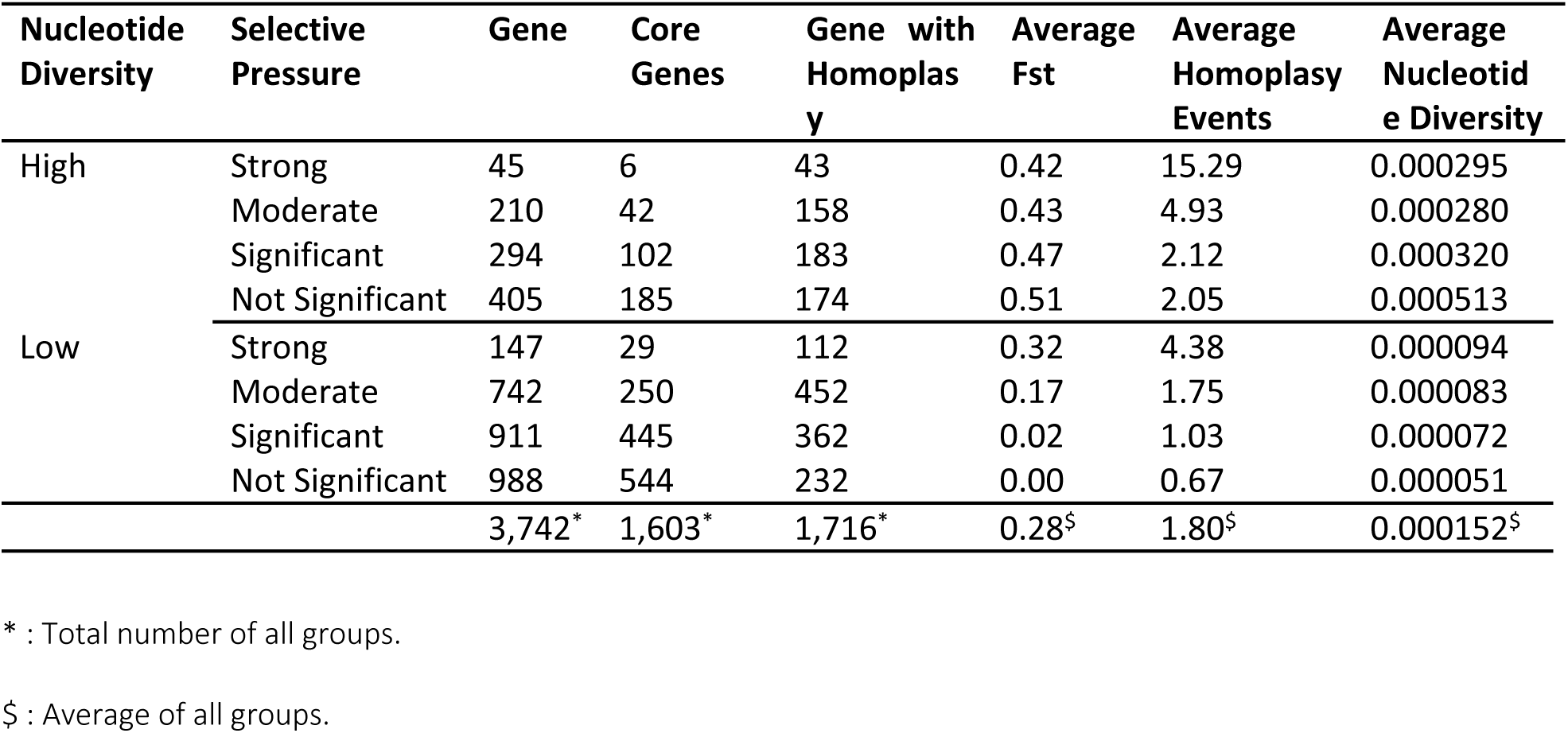
Population genetic characteristics under different selective pressures.

| Nucleotide Diversity | Selective Pressure | Gene | Core Genes | Gene with Homoplas y | Average Fst | Average Homoplas y Events | Average Nucleotide Diversity |
| --- | --- | --- | --- | --- | --- | --- | --- |
| High | Strong | 45 | 6 | 43 | 0.42 | 15.29 | 0.000295 |
|  | Moderate | 210 | 42 | 158 | 0.43 | 4.93 | 0.000280 |
|  | Significant | 294 | 102 | 183 | 0.47 | 2.12 | 0.000320 |
|  | Not Significant | 405 | 185 | 174 | 0.51 | 2.05 | 0.000513 |
| Low | Strong | 147 | 29 | 112 | 0.32 | 4.38 | 0.000094 |
|  | Moderate | 742 | 250 | 452 | 0.17 | 1.75 | 0.000083 |
|  | Significant | 911 | 445 | 362 | 0.02 | 1.03 | 0.000072 |
|  | Not Significant | 988 | 544 | 232 | 0.00 | 0.67 | 0.000051 |
|  |  | 3,742* | 1,603* | 1,716* | 0.28<sup>§</sup> | 1.80<sup>§</sup> | 0.000152<sup>§</sup> |

\* : Total number of all groups.
§ : Average of all groups.

When no signal of selective pressure was detected, 988 genes in the low nucleotide diversity group showed the lowest nucleotide diversity (average π = 5.11e-5), the lowest level of homoplasy (0.67 homoplasy events on average), and the lowest level of fixation between subpopulations (average Fst = 0), indicating extreme conservativeness in evolution of these genes in MTB. Another group of 405 genes in the high nucleotide diversity group showing no signal of selective pressure had the highest nucleotide diversity (average π = 5.13e-4) and also the highest level of fixation between subpopulations (average Fst = 0.51) and a moderate level of homoplasy (2.05 homoplasy events on average), indicating that population stratification was the major contributor to the nucleotide diversity for this group and a relatively low level of conservativeness in evolution for these genes. The proportion of core genes showing no signal of selective pressure in the group of high nucleotide diversity (185/405, 45.68%) and the group of low nucleotide diversity (544/988. 55.06%) were higher than the overall level (1,603/3,742, 42.84%, P-value < 0.001); the group of low nucleotide diversity was significantly higher than the group of high nucleotide diversity as well (P-value < 0.001). As mentioned above, there were 40 core genes showing no genetic variation, which made the conserved core genes to be 584 in total. Genes involved in translation (GO:0006412) and regulation of DNA-templated transcription (GO:0006355) were over-presented in this group by 43 and 50 genes, respectively. (**Table 2**)

## Discussion

Our study has shown that MTB has a highly conserved pangenome with only 85 new genes not present in the reference genome H37Rv in a diversified sample set. The size of pangenome of MTB is only one-fifth of *E. coli*, but the proportion of core genes is three times of that in *E. coli* at protein level and eight times at gene level.^51^ This small pangenome size is presumably decided by the obligate intracellular lifestyle of MTB.^52^ The average nucleotide diversity for coding genes is at least one order lower than that in other organisms.^53,54^ These findings agree with and further confirm the notion that MTB is evolutionarily conserved as reported by previous studies using a standard reference genome or focusing on a specific region.^55–57^

Due to interruptions of ORFs caused by various types of genetic variation, about half of the core coding genes were potentially dysfunctional in different genomes even with the presence of CDSs. Structural variation accounts for more than half of the observed interruptions in coding genes while high impact SNPs only correspond to about ten percent, suggesting SVs are a major source of genetic variations and is the major contributor to adaptive loss in evolution.^58^ Though genetic variation due to LSPs are relatively limited in MTB, we have identified two genes, *pimC* and *suoX*, which are deleted due to the LSP of RvD2, may have crucial impact on the metabolism. PimC is an amannosyltransferase involved in cell wall biosynthesis and some of its products are important virulence factors in MTB.^59,60^ Since conserved domain searching hasn’t identified any other homologues gene with similar function in the pangenome (data not shown), we assume there is an alternative pathway involved in the biosynthesis of Ac_n_PIM_3_ in Lineage 2 which has not yet been discovered. *SuoX* oxidizes sulfite to sulfate directly in sulfur metabolism. Due to its nucleophilicity and strong reductive capacity, sulfite can be toxic for bacteria but can be detoxified by oxidation to sulfate.^61^ Direct oxidation of sulfite seems to be the exclusive way to detoxify sulfite via oxidation, yet the gene *suoX* encoding sulfite oxidase in MTB is deleted in Lineage 2 strains and no other sulfite oxidase has been identified in MTB so far, which means the direct oxidation of sulfite in Lineage 2 may be compromised.^61^ In the reverse direction, the assimilation pathway of sulfate, sulfite could be further reduced to sulfide, which is the form of sulfur required for biosynthesis of sulfur containing metabolites such as cysteine and acetate.^62^ Sulfate assimilation is important for bacteria persistence. For example, H_2_S has been shown to stimulate respiration, growth and pathogenesis of MTB *in vivo*.^63^ MTB can produce endogenous H_2_S from cysteine by a desulfhydrase (probably *Rv3684*).^64^ The absence of *suoX* in Lineage 2 might contribute to the higher virulence of Beijing strains observed, as compromised oxidation capacity would increase the concentration of sulfite, which in turn moves the balance towards the direction of assimilation and consequently increases the production of H_2_S. These observations indicate that although LSPs only have a limited contribution to the difference in genetic content, some of these differences may have important consequences that their absences in certain genotype could result in differentiation in metabolism that might have contributed to the prevalence of certain genotypes.

The landscape of nucleotide diversity for individual genes suggests that different genes in MTB undergo very different evolutionary processes, which may be influenced by several factors. We have identified 2,349 out of 3,742 coding genes under significant selective pressure in MTB which is very close to the number (2,729 out of 3,979) reported in another study using dN/dS ratio as index, indicating the robustness of both methods.^65^ The number of genes located in cell wall and membrane structures under strong selective pressure is almost doubles that in plasm, which might reflect the intracellular parasite lifestyle with limited sources of nutrients and constant attacks from host immune system, as most of these processes are taking place at the cell wall and membrane region. Selective pressure alone could explain about 30% of the nucleotide diversity observed in our dataset, and its strength is positively related to the intensity of positive selection in spite of other factors, indicating though it may be slow, the conservative MTB genome is still evolving. One important direction in evolution might be the interaction between MTB and the host immune system. Genes involved in disruption of host structures are over-presented in the group with high level of positive selection, indicating these selected mutations might provide advantages in fighting host immune attacks. Especially metabolism of fatty acid may have played a key role as indicated by the over-presentation of genes with phospholipase activity and in fatty acid biosynthesis process.

The population structure of MTB also has a heavy influence on the landscape of genetic diversity, which has been addressed in genome wide association studies.^66^ Because there is no horizontal gene transfer or random mating in modern MTBC strains, any genotype or a random strain and all its descents could be considered as a clonal subpopulation.^67^ Therefore, population stratification, defined as systematic differences between subpopulations in allele frequences, is expected to be high in MTB. Indeed, we have detected various levels of population stratification in groups of genes undergoing different evolution processes. However, population stratification is missing in a group of conserved core genes. In addition, this group of genes also has the lowest average nucleotide diversity and the lowest level of homoplasy and thus are evolutionarily conservative, indicating strong background selection working here. Background selection, or purifying selection, has been demonstrated as G/C-skew in MTB and been proposed as the base line for genetic variation.^68,69^ Our data confirmed that the majority of coding Rv genes have significant positive G/C-skew (data not shown) which is expected for genes under background selection. Background selection constantly removes deleterious mutations in the population by natural selection, which reduces neutral diversity and is supported by the observed low nucleotide diversity in MTB. However, the strength of background selection seems to vary between different biological processes, as some fundamental processes, such as translation, regulation of transcription and regulation of growth, are over-presented in the most conserved gene group, which can be expected.

To summarize, while population stratification introduces systematic differences and thus increases genetic variation, background selection decreases diversity; selective pressure works as a resistive force to these two factors through ways such as positive selection. Selective pressure reduces systematic genetic variation introduced by population stratification and increase the frequency of emerging advantageous mutations and thus genetic variations against background selection. This combination of complex interactions explains the observed divergent patterns of nucleotide diversity for genes under no selective pressure, which represents genes predominantly under the influence of background selection or population stratification; when selective pressure is present, nucleotide diversity converges to the average level as the strength of selective pressure increases.

A few exceptions were observed. One example is the *rpsL* gene, which shows a high level of homoplasy (104 homoplasy events) indicating positive selection of advantageous mutations, but no signal of selective pressure. Mutations in codon 43 in *rpsL* confer a high-level resistance to streptomycin. Streptomycin is the first anti-TB drug used in monotherapy of tuberculosis but has been replaced by oral drugs for almost two decades now.^70^ The disappearance of signal of selective pressure reflects the changes in the standard treatment regime. Though the historical selective pressure no longer exists for *rpsL* it could still be detected with signal of positive selection. Another factor that might have contributed to the persistent signal of positive selection is fitness cost. Mutations in other drug resistance related genes, such as *rpoB*, usually cause different levels of fitness cost.^71^ While strains with mutations in *rpsL* conferring streptomycin resistance are not affected by such deficiency.^72^ Epistatis may also play a role in the high frequency of *rpsL* mutantions as shown in a study with *E.coli* which reported higher fitness in some double mutants of *rpoB/rpsL* than wild genotypes.^73^ Also in our previous analysis using the same sample set, the association between drug resistance patterns and genotypes suggests that certain genotypes which developed resistance to streptomycin seem not to show deficiency in further developing rifampin resistance.^74^ However, this type of highly advantageous mutations without fitness cost in genes is very rare, as we only observed 32 genes in this category with no signal of selective pressures and a high level of homoplasy. Further checking the function of these genes might be helpful to identify historical selective pressure in the evolution of MTB. These observations suggest that the current landscape of genetic diversity in MTB is the outcome of complicated interactions between changing selective pressures, background selection, clonal population structure, and other factors such as fitness cost of mutations.

Our study has shown the necessity and advantages to switch from an approach using a standard reference genome focusing on SNPs and small indels to a more comprehensive pangenome approach with detailed genetic variation information. By combining pangenome, population genetics and gene ontology, new insights into the biology and the interaction of MTB with its environment may be gained. Future studies with more advanced sequencing techniques like long reads sequencing and more versatile bioinformatic tools such as graph pangenome will remove the drawbacks in current methodology and improve our understanding about the epidemiology and pathogenicity of MTB.^75^

## Funding

National Key Research and Development Project (2022YFC2305204)

## Notes

### Competing Interest Statement

The authors have declared no competing interest.

